# C/VDdb: a multi-omics expression profiling database for a knowledge-driven approach in cardiovascular disease (CVD)

**DOI:** 10.1101/430660

**Authors:** Marco Fernandes, Alisha Patel, Holger Husi

## Abstract

The cardiovascular disease (C/VD) database is an integrated and clustered information resource that covers multi-omic studies (microRNA, genomics, proteomics and metabolomics) of cardiovascular-related traits with special emphasis on coronary artery disease (CAD). This resource was built by mining existing literature and public databases and thereafter manual biocuration was performed. To enable integration of omic data from distinct platforms and species, a specific ontology was applied to tie together and harmonise multi-level omic studies based on gene and protein clusters (CluSO) and mapping of orthologous genes (OMAP) across species.

CAD continues to be a leading cause of death in the population worldwide, and it is generally thought to be an age-related disease. However, CAD incidence rates are now known to be highly influenced by environmental factors and interactions, in addition to genetic determinants. With the complexity of CAD aetiology, there is a difficulty in research studies to elucidate general elements compared to other cardiovascular diseases.

Data from 92 studies, covering 13945 molecular entries (4353 unique molecules) is described, including data descriptors for experimental setup, study design, discovery-validation sample size and associated fold-changes of the differentially expressed molecular features (p-value<0.05). A dedicated interactive web interface, equipped with a multi-parametric search engine, data export and indexing menus are provided for a user-accessible browsing experience.

The main aim of this work was the development of a data repository linking clinical information and molecular differential expression in several CVD-related traits from multi-omics studies (genomics, transcriptomics, proteomics and metabolomics). As an example case of how to query and identify data sets within the database framework and concomitantly demonstrate the database utility, we queried CAD-associated studies and performed a systems-level integrative analysis.

URL: www.padb.org/cvd

## Introduction

Cardiovascular disease (CVD) encompasses many pathological conditions that lead to an integral loss in function of the heart, cardiovascular structures and the circulatory system. This life-threating and highly debilitating condition is responsible for 32% of all total global deaths, of which more than half,17%, are due to ischemic heart disease (IHD), 10% stroke and nearly 2% due to hypertensive heart disease (HHD) [1].

Atherosclerosis is the leading underlying condition for the establishment of CAD and the predominant cause of IHD that often leads to myocardial infarction (MI) [2, 3]. CAD in the clinical manifestation of atherosclerosis refers to chronic inflammation with early and gradual accumulation over the life-span of lipids and fibrous elements, that form the basis of the atherosclerotic plaque in large and medium-size arteries essential in the transport of oxygen and nutrients to the heart [2, 4, 5]. In most of the cases the disease remains clinically asymptomatic during the first decades of its development [6], until the surface of the atherosclerotic plaque suffers rupture or erosion, causing formation of a thrombus that obstructs the vessel with deprivation of an adequate blood-flow and consequent ischaemic tissue damage [7]. This ultimately leads to severe disease manifestations as myocardial infarction (MI), stroke, or acute coronary disease (ACD) [8]. CAD is perceived as a complex disease of multifactorial origin with a strong genetic component and with an estimated familial heritability between 30 to 60% [9–11]. Despite major improvements in many of the modifiable risk factors, such as smoking habits, alcohol intake, and total cholesterol amongst Western nations over the years [12], CAD still remains the main cause for the majority of cardiovascular events [1].

Several low-income countries have started to adopt into their culture the Western-lifestyle, and with it many of the CVD-modifiable cardio-metabolic risk factors, and therefore is expected that this will certainly elevate the rate and range of CVD epidemiology globally [13, 14]. The total annual expenditure of CVD, adding-up direct and indirect costs across EU countries ascends to 210 billion euros, [12] likewise CVD and stroke altogether have an annual impact over the United States economy of up to 258 billion euros [15].

In the last decade the emergence of high-throughput Omics platforms opened the possibility to mechanistically understand diseases and diseases stages at the molecular level. These analytic technologies are able to provide reliable, robust and fast detection of candidate biomarkers for disease onset and progression [16].

Screening of tissues and body accessible fluid sources for the detection of small (~21-mer), highly conserved non-coding RNA molecules - miRNAs, that regulate post-transcriptionally the expression of genes by binding based on sequence complementarity to the 3′-untranslated regions (3′-UTR) of specific mRNAs [17] and exert degradation or translational repression of mRNA targets [18], provides another layer of understating over the role and how they confer robustness to transcriptional programs in cell differentiation, proliferation, survival and in disease [19].

On the other hand, transcriptomics technologies are of invaluable source by capturing patterns of expression of ribonucleic acid (RNA) transcripts across the genome [20]. Then, assessing gene expression through microarrays, deep sequencing by RNA sequencing (RNA-seq) [21, 22] and with other complementary whole-genome sequencing platforms [23] offers a way to identify signature genes related to a certain human condition [21]. Moreover, the study of the proteome enables the characterization of the potential function of genes asserted by proteins participating in molecular signalling cascades, as catalysts of enzymatic reactions, and in protein-protein physical interactions within many cellular processes in health and disease [20]. Therefore, detection and quantitation using mass spectrometry (MS)-based proteomics and combined with traditional gel-based to liquid chromatography (LC) coupled with high-resolution tandem mass spectrometry (MS/MS) has become a widely-accepted method for measurement of proteins. [24, 25].

Metabolites are not only the end-products of enzymatic reactions, but also active participants in the regulation of the cellular microenvironment, in which changes in abundance at the intracellular level of metabolite intermediates can promote changes in gene signal transduction [26, 27] and therefore can be used to investigate the general health status [28].

Databases handling high-throughput expression profiles of miRNAs, genes, proteins and metabolites in pathological conditions and disease stages are still scarce, exceptionally in the case of general scope transcriptomics repositories such as NCBI Gene Expression Omnibus (GEO) [29] and EBI Expression Atlas [30].

However, the core source for publicly available omic data still resides in peer-reviewed publications, therefore we developed a clustered molecular repository for cardiovascular and vascular diseases, C/VD, containing expression profiles of microRNA, protein and metabolite across tissues, body-accessible fluids, and species derived of collected experimental data from peer-reviewed research publications and other relevant databases. Additionally, we performed as a show case of the database utility and to illustrate potential uses of the data enclosed in our newly established database, a meta-analysis integrating omics data sets at the pathway level in CAD.

## Material and methods

### C/VD database construction

#### Data collection, storage and handling

The content of the C/VD database is based on manual extraction of data from available literature on the topic of CVD and omics technologies. We used several search strings in PubMed and Scopus in order to cover the topic as much as possible (e.g. “Cardiovascular Diseases”[Mesh] AND (miRNA OR genomic* OR proteomic* OR metabolomic*) NOT review) and filtered to display only studies in English language. Additional gene and miRNA data sets were retrieved from the Gene Expression Omnibus (GEO) Database [29]. We kept data in the tabular form and retained information of the original source for easy data tracking.

#### Data biocuration

Publication-based molecular identifiers were converted to internal identifiers of the Pan-omics Analysis Database (PADB) initiative (www.padb.org), which uses the clustered sequences and orthologs (CluSO) identifiers and an ortholog mapping resource (OMAP). These internal identifiers allow us to reduce data redundancy found in many multi-omic studies. Moreover, it ensures that the C/VD database remains fully integrated into other databases held within the PADB infrastructure.

We only retained statistically significant and differently expressed molecules e.g. raw and/or adjusted for multi-comparisons p-value <0.05, and fold-change (FC), transcriptomics: |FC|≥ 2.0, proteomics and metabolomics: |FC|≥ 1.3.

#### Database deployment

C/VD relies on a NoSQL database system based on dynamically interlinked and pre-assembled HTML files using in-house built software as described elsewhere in more detail [31]. Briefly, the generated database relies on a NoSQL-system, whereby source data is manipulated in spreadsheets and parsed into pre-assembled and interlinked html files. In C/VD there are three table sources handling data concerning demographics (clinical data), experimental setup descriptors and molecular descriptors. All of these tables follow the standard database nomenclature and structure, i.e. they are all linked to each other and to external databases, for instance UniProt/SwissProt [32], ChEBI [33], Entrez GeneID [34], miRbase [35], Ensembl [36], UniGene [34] and the Online Mendelian Inheritance in Man (OMIM) database [37]. Thus, in this way the customised parser allows to emulate a database system by pre-assembling query outputs by for example grouping based on disease/ tissue/ molecule.

Conversion of the collected data from source tables to html pages that constitute the front-end of the database was ensured via in-house built software. Interactive controls were also added to the html tables such as sorting, pagination, multi-column sorting and data export features (Excel, PDF, CSV) by incorporating the DataTables plugin [38] for the jQuery JavaScript library [39].

A search interface was also implemented in the database that relies on the Hypertext Pre-processor (PHP) [40] scripting language, based on summary tables i.e. rearranged tables that simplify the underlying queries, therefore allowing fast, enhanced and customised searches across the database.

#### Availability and requirements

Home page: Accessible through the PADB portal at http://www.padb.org/cvddb. Operating system: C/VD is a web-based database thus it is platform independent. Requirements: works in any browser supporting HTML 4.01 and Java. License: free for academic use, but requires a license for any commercial purposes.

### Case study with a C/VD subset: CAD

#### Data selection, pre-processing and meta-analysis

We selected a subset of C/VD, CAD data sets of human cohorts, screened in blood as comparison between CAD cases and control groups in at least two omics technologies. Since, C/VD only handles data below the *P*-value cut-off of 0.05, redundant molecular entries within the same study were combined by averaging their expression values and p-values were combined based on the calculation and combination of intra-studies *P*-values by using the popular and well-established Fisher’s method [41]. Other methods exist, but Fisher’s method performs better in our data type setting, since Stouffer’s method Stouffer [42] requires studies with similar quality, such as sharing the same detection platform and also requires similar number of missing values.

A vote counting strategy [43] was used to combine molecular information, where differentially expressed (DE) molecules are first selected based on an overall statistical—*P*-value threshold defined as *P* <0.05. Then, dependent on the type of - omics technology/ platform, different fold-change (FC) cut-offs were applied: genomics, including miRNA and gene expression |FC|≥2.0; proteomics ≥1.5 and metabolomics |FC|≥1.3 to obtain a list of DE molecules for each study. The vote for each molecule can be then calculated as the total number of times (frequency (%) distribution) it occurred in all DE study lists to be combined. Afterwards, the final DE molecules list can be selected based upon a set of minimal number of votes. This simple approach and yet robust for further downstream analysis, outperforms other meta-analysis methods, since most cannot handle highly heterogeneous data types with a high degree of missing values. Then, based on the frequency distribution of the molecular regulation directionality (down or up-regulation trends) across three fold-change (FC) cut-offs (|FC|≥1.3, 1.5 and 2.0) observed in microRNA (MIR), gene (mRNA), protein (PRO) and metabolite (MET) expression data. Prevalence was given to any molecular entity observing frequency ≥60% in any case of contradictory regulation.

#### Data dimensionality reduction

Principal component analysis (PCA) and non-metric multi-dimensional scaling (MDS) analysis were performed in Primer-E (version 6.1.6) [44]. The input data for the PCA was a matrix containing differential expression as log_2_ fold-change (log_2_FC) of each molecular entity (as variables, N=654; miRNA, protein and metabolite) across several Omics study types (as samples, N=75). In this analysis we used molecular data reported in at least three independent studies; handling of missing data entries was resolved by setting it to zero. The triangular resemblance matrix required for MDS was based on the measure of Euclidean distances. MDS subsets were also generated to visualise in the 2D and 3D space superimposed Omics studies.

#### Functional enrichment and pathway analysis

The most reported molecular entries of each type of omic technology were the input for term clustering analysis performed using gene ontology (GO) annotation including biological process (BP), molecular function (MF) and cellular component (CC); pathway term clustering using the Kyoto encyclopedia of genes and genomes (KEGG) [45] and WikiPathways [46] annotation built-in ClueGO app [47] for the Cytoscape [48] network analysis platform. The default parameters were used (or otherwise in-text specified) with an overall statistical significance value set to *P*-value <0.05.

#### Supervised development of a multi-omics integrative molecular model

The development of a molecular model in CAD followed an iterative design (Fig 1) involving data collecting, establishment of a multi-omics database, C/VD, meta-analysis, molecular clustering into GO and pathway terms and analysis of the interactome followed by several iteration swaps until the finalised output model was reached.

**Fig 1.**
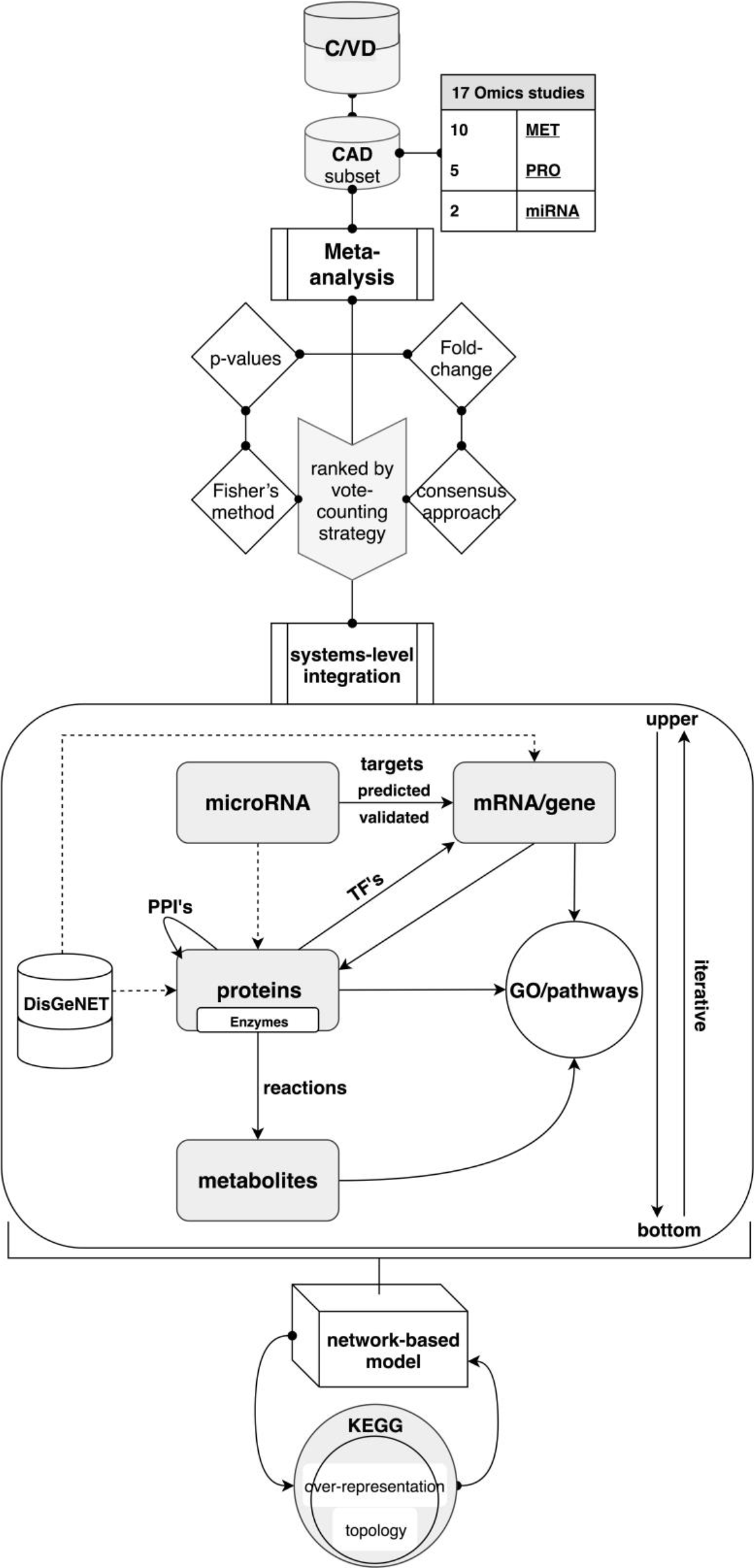
Supervised development of an integrative multi-omics molecular model in CAD. Coronary artery disease (CAD) multi-omics data sets retrieved from C/VD regarding molecular entity type, for instance metabolite (MET), proteins (PRO) and microRNA (miRNA). Molecular entities from the studies were initially ranked by meta-analysis using a vote-counting approach (frequency assessment), in which statistical scores (p-values) were combined by implementing Fisher’s method and differential expression of molecules (fold-change) were merged through a consensus approach. Following the meta-analysis, a combinatorial integration at the systems-level of all molecular entities concerning modulation of transcriptional activity via assessment of transcription factors (TF’s), post-transcriptional regulation ascertained by microRNA-gene interactions (computationally predicted and experimentally validated), protein-protein interactions (PPI’s), evaluation of metabolic flux and regulation via association of gene-enzyme-reaction-compound and over-representation analysis of gene ontology (GO) terms and KEGG pathways. Further contextualisation of the disease model was ensured via incorporation of known CAD-gene associations sourced from the DisGeNET [44] repository.

Post-transcriptional regulation by microRNAs (miRNAs) was verified by assigning miRNAs-targets interactions using pre-computed target predictions (miRanda-v5 update of 19-07-2012) [49] and validated (mirTarBase update of 15-06-2016) [50] mRNA/gene targets via the CluePedia app 1.5.0 [51] and complementary single searches on an *ad hoc* basis in mirPath v3.0 [52].

Assessment of transcriptional regulation by transcription factors (TF’s) was ensured by the use of CyTargetLinker 3.0.1 [53] and GeneMANIA 3.4.1 [54]. Both apps work within the Cytoscape network analysis platform [48].

Protein-protein physical interactions (PPI’s) were assessed using STRING 1.3.2 [55] by establishing an overall minimum confidence score of the enrichment to 0.7.

Enzymatic reactions linking metabolites with proteins were derived from the MetScape app 3.1.3 [56] and missing matches were completed with manual searches across KEGG [45] and Brenda databases [57] on an *ad hoc* basis.

In each omic layer (miRNA, protein and metabolites) a subroutine concerning convergence to gene ontology (GO), KEGG [45] or WikiPathways [46] pathway terms was created to ensure a focus and validity of the model.

Contextualisation of the created model regarding the studied disease, CAD, was ensured by adding the highest scored gene-disease associations (GDA) from the DisGeNET database [58] and reconnecting all the molecular interactions.

At the end, the calculated overall fold-changes of all molecules derived from the meta-analysis were incorporated and colour coded on top of the built network-based model.

The molecular content of the developed model was then tested for over-representation (hypergeometric test) and pathway topology analysis using KEGG [45] pathway terms from the web interface of MetaboAnalyst [59].

## Results

### Database structure and navigation

Navigation through the C/VD database front-end can be either by browsing the study, sample/demographic, tissue/source or by molecular entry index pages, or alternatively through molecule names, tissue or disease queries in the search interface (Supporting information Figs S1-S5).

#### Statistics and data summary of C/VD

The database consists of 13945 molecule entries (redundant) of which 4353 are unique (non-redundant) from a collection of 98 manually curated studies of 50 publications related with CVD. A summary of the database content using a snippet of the categorical and numeric variables (Fig 2) allows uncovering data trends amongst the many datasets that C/VD comprehends by plotting a histogram-like chart built using the Tableplot R-package [60]. From observation of the tableplot, the main molecular entries populating the C/VD database corresponds to human, detected in blood, mainly granulocytes and mononuclear cells from mRNA/gene in coronary heart disease (CHD). On the other hand, amongst all other molecular entities in C/VD, microRNAs (miRNA) tend to have lower raw p-values and higher fold-change (FC) values. Additionally, studies handling metabolomics profiling (MET) tend to have larger number of individuals displayed here as cohort/sample size (N) (Fig 2).

**Fig 2.**
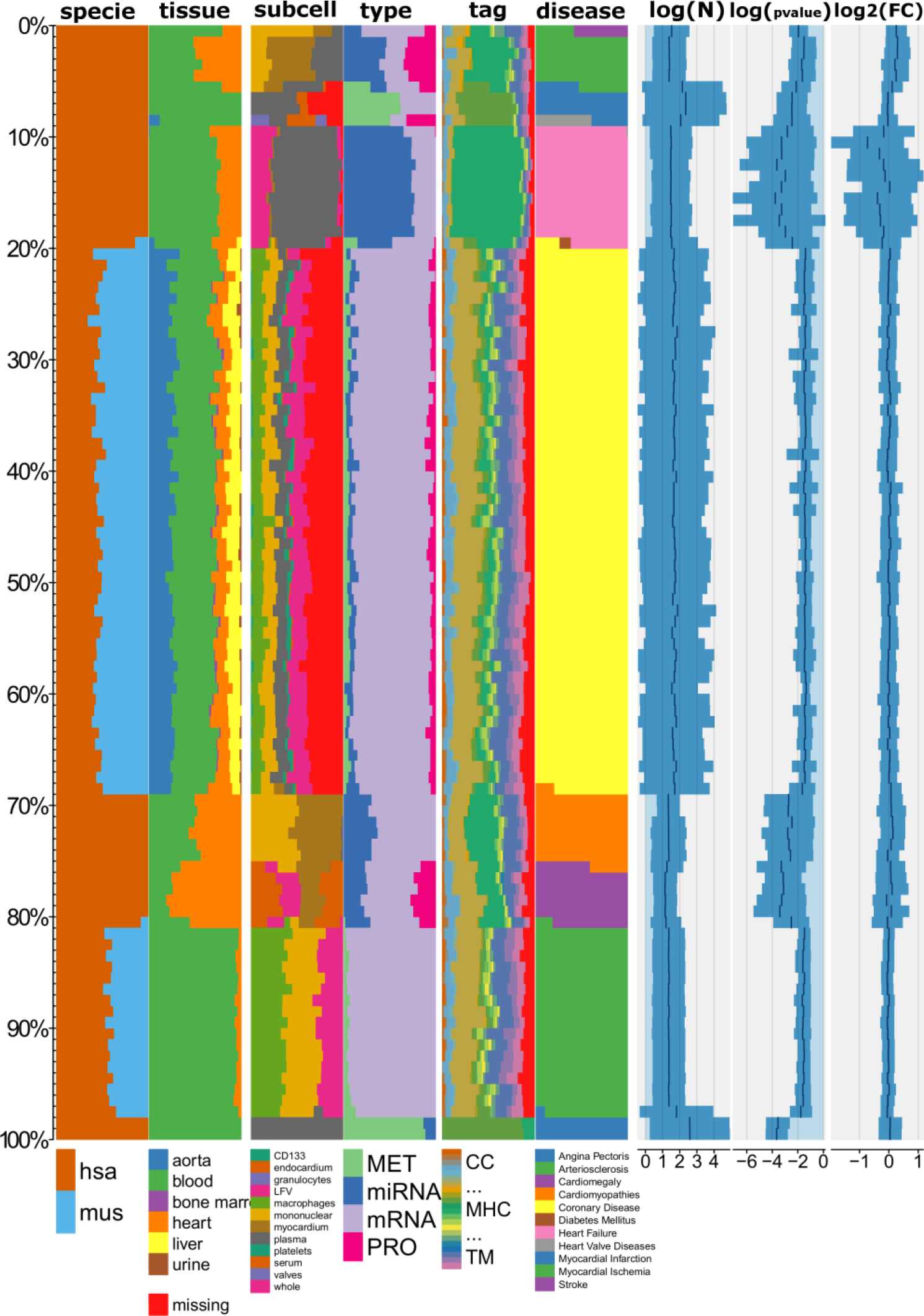
Tableplot of the C/VD database. Each column represents a variable and each row (bar) is an aggregate of a fixed number of records, i.e. a row bin (here equivalent to a molecule entry). The numeric variables (log10 (N), log10 (p-value) and log2 (FC)) are displayed as bar charts and categorical variables (specie-disease) as stacked percentage bar charts. Disease type was used as the sorting variable. Cohort size (N), log10(p-value), log2(fold-change(FC)). Species: *Homo sapiens* (hsa), *Mus musculus* (mus), Tags: ENZ: enzyme, DIS: disease, SIG: signalling, INH: inhibitor TF: transcription and translation, gene regulation, TP: transport, storage, endocytosis, exocytosis, vesicles, CS: Cell shape, MOD: modulator, regulator, MET: metabolite, TM: transmembrane, CHA: chaperone, chaperonin, MIR: microRNA, SNORNA: small nucleolar RNA, DEV: development, cell growth, differentiation, morphogenesis, CC: cell cycle, RIB: ribosome, CNL: channel, RCP: receptor, IGG: Immunoglobulin, SCA: scaffolder, docking, adaptor, MHC: major histocompatibility complex component.

Almost half (48.53%) of the C/VD content (Fig 3A) within a multitude of omics studies concerns coronary heart disease (CHD), followed by arteriosclerosis, 16.89%, and heart failure, 11.00%, amongst other conditions that fall under the 10% representation within C/VD. Most conditions are characterised at least by two types of omics technologies, with the exception of calcinosis, diabetes mellitus, cardiac resynchronization therapy (CRsyT) and stroke (Fig 3A).

**Fig 3.**
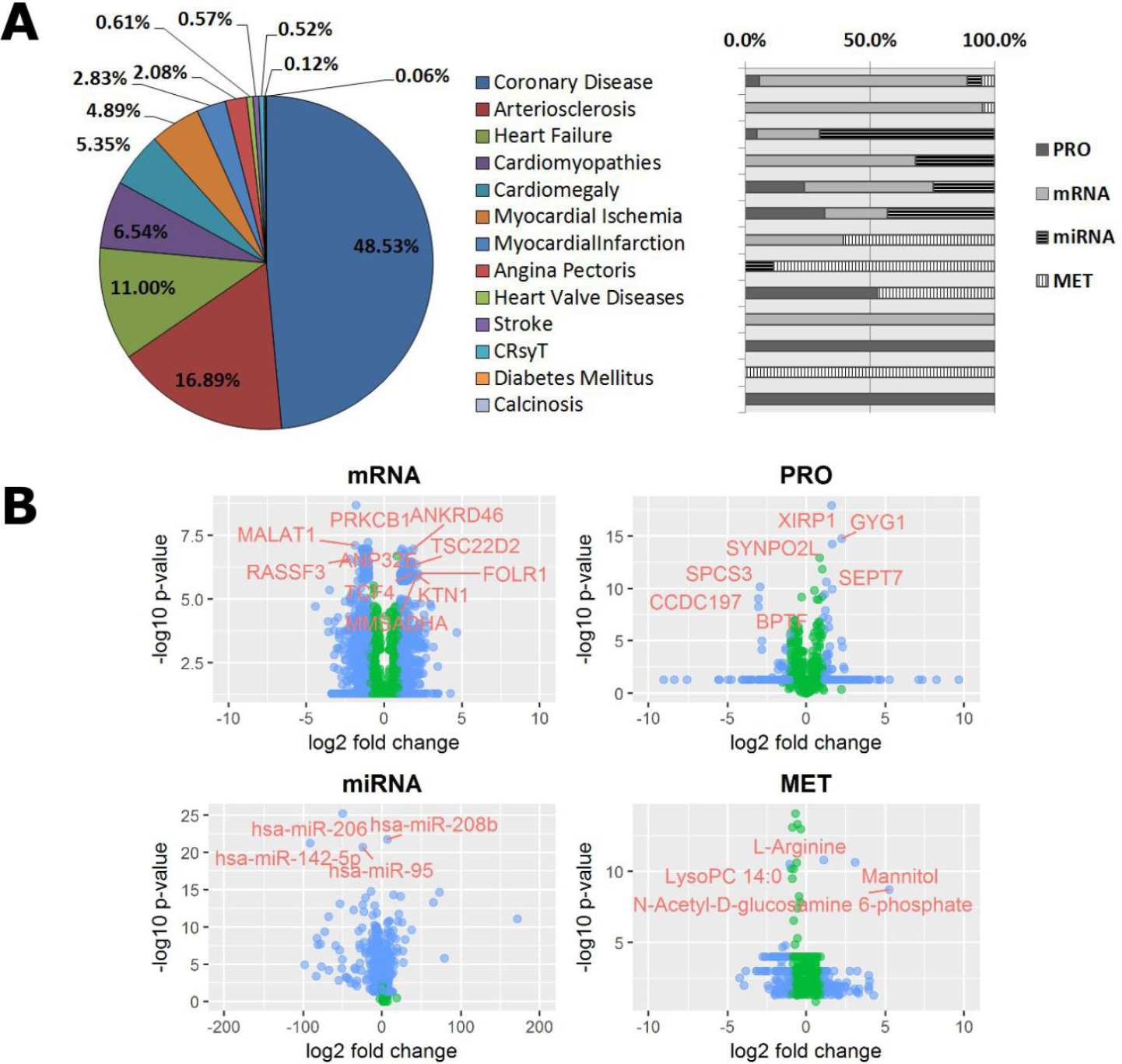
Disease representativity in the C/VD database (A) and volcano plots displaying differential expression in each molecular entity per type of Omics technology (B). IC10 (2016) disease classification of the content of C/VD database and relative proportion (%) to Omics studies. Proteomics (PRO), mRNA (gene-array), miRNA (miRNA-array), and metabolomics (MET). Cardiac Resynchronization Therapy (CRsyT). Each variable pair shows differential expression across each molecular entity, X-axis represents log2(fold-change) and Y-axis, −log10(p-value). Type of Omics: Proteomics (PRO), mRNA (gene-array), miRNA (miRNA-array), and metabolomics (MET). The blue filled data points denotes molecular entities than observe the following thresholds: mRNA, (Log2FC)>1.8 & Log10pvalue>5.5; miRNA, (Log2FC)>1.8 & Log10pvalue>17; PRO, (Log2FC)>1.5 & Log10pvalue>5 and MET, (Log2FC)>1.8 & Log10pvalue>5, and the green filled data points the ones than did not observe that condition. mRNA: Metastasis associated lung adenocarcinoma transcript 1 protein (MALAT1), RAS association domain-containing protein 3 (RASSF3), Protein kinase C, beta type 1 (PRKCB1), Ankyrin repeat containing protein 46 (ANKRD46), TSC22-related inducible leucine zipper 4 (TSC22D2), Folate receptor alpha precursor (FOLR1), Kinectin (KTN1), Methylmalonate-semialdehyde dehydrogenase [acylating] (MMSADHA), Transcription factor 4 (TCF4), Acidic (leucine-rich) nuclear phosphoprotein 32 family member E (ANP32E). PRO: XIN actin-binding repeat-containing protein 1 (XIRP1), Glycogenin-1 (GYG1), Synaptopodin 2-like protein (SYNPO2L), Signal peptidase complex subunit 3 (SPCS3), Spectrin repeat containing protein (CCDC197), Septin-7 (SEPT7) and Nucleosome-remodelling factor subunit BPTF (BPTF).

Representation of each omics technology type from C/VD by plotting molecular expression and statistical scores (Fig 3B) helps in the first instance to assess data dispersion/variability and outlier detection. Here, it can be verified that many data points in proteomics (PRO) and metabolomics (MET) display a linear rearrangement perpendicular to the y-axis, which corresponds to the overall statistical cut-off selected by the authors in many experimental omics studies. On the other hand, gene (mRNA) and microRNA (miRNA) differential expression studies tend to have associated for each molecular entry individual statistical scores, instead of an overall cut-off (Fig 3B).

The displayed gene/proteins (Fig 3B) with higher fold-change and p-value thresholds such as RASSF3, XIRP1, SYNPO2L and SEPT7 are involved in cell shape remodelling, PRKCB1, MMSADHA, GYG1 and SPCS3 have enzymatic properties, TCF4 and BPTF have roles in transcription and translation (gene regulation), ANKRD46 and KTN1 are transmembrane proteins functioning as gateways of many substances, and FOLR1 is an anchor element in biological membranes acting as receptor. Others like MALAT1, TSC22D2 and CCDC197 have unknown roles. Highly expressed metabolites such as the LysoPC(14:0), a lysophospholipid with roles in lipid signalling and fatty acid (FA) metabolism; L-arginine an essential amino acid (AA) with numerous functions within the body; mannitol and N-Acetyl-D-Glucosamine 6-Phosphate (GlcNAc-6-P) are sugars based endogenous metabolites, the first an sugar alcohol with diuretic functions and the latter is an acylaminosugar. Both are involved in the phosphotransferase system (PTS) in bacteria what could suggest an association of cardiovascular disease with metabolites derived from the intestinal microbiota. Regarding microRNAs expression, hsa-miR-95, hsa-miR-142-5p and hsa-miR-206 have as predicted targets, genes involved in proteolytic cascades mediated by ubiquitin. Similarly, hsa-miR-208b targets genes with functional roles in the p53 signalling pathway (Fig 3B).

#### Data dimensionality reduction

The application of data dimension reduction methods (Fig 4), with principal components analysis (PCA) (Fig 4a) that describes 93.9% of the cumulative variance of PC1 (75.5%) and PC2 (18.4%) generated from the content of the C/VD database based on a matrix handling differential expression of each molecular entity per Omics type as variables and study ID (“Exp”) as samples. The PCA plot shows the score of each disease case and the loadings of each variable i.e. the molecular elements on the first two principal components. The greater spatial distance among most of the disease cases that overlap in the plot and heart failure (HF) suggests that the latter can be an outlier due mostly to contribution of microRNA expression, with hsa-miR-208b (BZJ12), hsa-miR-208a (BZJ35), hsa-miR-125b-1-3p (BZQ81) and hsa-miR-376a-2-5p (BZ615) as main molecular culprits. The visualisation of the level of resemblance among many CVD disease cases of the highly overlapped cluster (Fig 4a) by a non-metric multi-dimensional scaling (MDS) method (Fig 4b1, 4b2) made possible to pairwise compare spatial distances at 2D (Fig 4b1) and 3D (Fig 4b2) among the origin of tissue/ fluid sources, e.g. blood and heart tissue.

**Fig 4.**
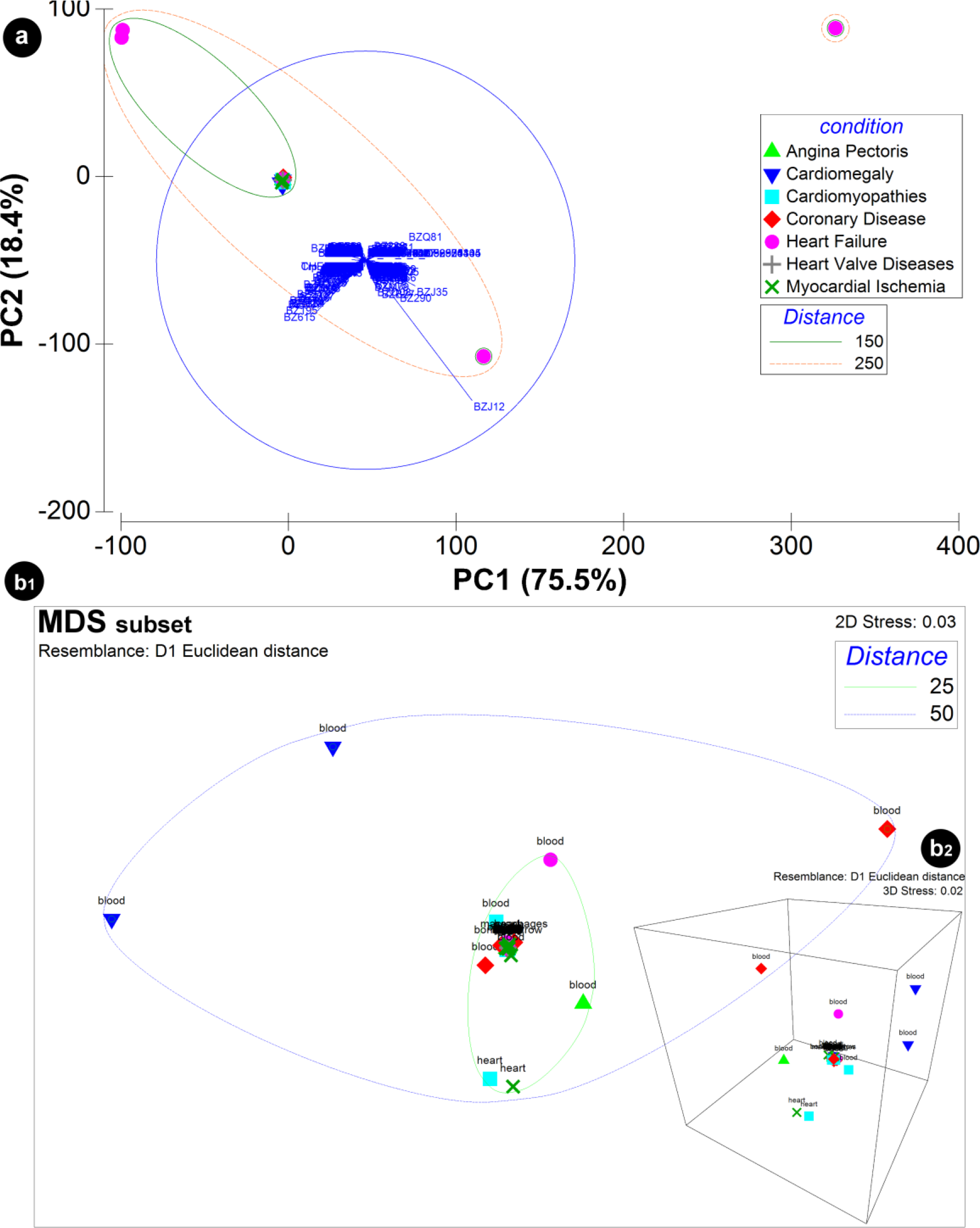
Principal components analysis (PCA) of the content of the C/VD database. Data input was a matrix with differential expression of each molecular entity per Omics type as variables and study ID (“Exp”) as samples. IC10 (2016) disease classification was used as categorical labels. The PCA (a) displays both the PCA scores, X-axis: PC1 (75.5%), Y-axis: PC2 (18.4%) with the confidence ellipse of 95% surrounding CVD conditions and the PCA loadings with the molecular variables hsa-miR-208b (BZJ12), hsa-miR-208a (BZJ35), hsa-miR-206 (BZ290), hsa-miR-125b-1-3p (BZQ81) and hsa-miR-376a-2-5p (BZ615) represented. Visualisation of the level of similarity between CVD conditions of the highly superimposed cluster (a) was achieved by implementing a non-metric multi-dimensional scaling (MDS) approach in the 2D (b1) and 3D (b2) space. The MDS plot exhibits pairwise dissimilarities as a measure of the Euclidean distances among data points. Tissue/ fluid sources (blood, heart) are represented as a factor of the MDS analysis.

### Case study with a C/VD subset: CAD

#### Data sets description

We selected a subset of 21 studies from the C/VD database regarding CAD across miRNA (two studies), protein (nine studies) and metabolite (10 studies) expression detected in blood of human cohorts (Table 1). Additional description of other clinical parameters such as age, gender, and clinical history of cases and controls is available in Supplementary Table S1.

**Table 1.**
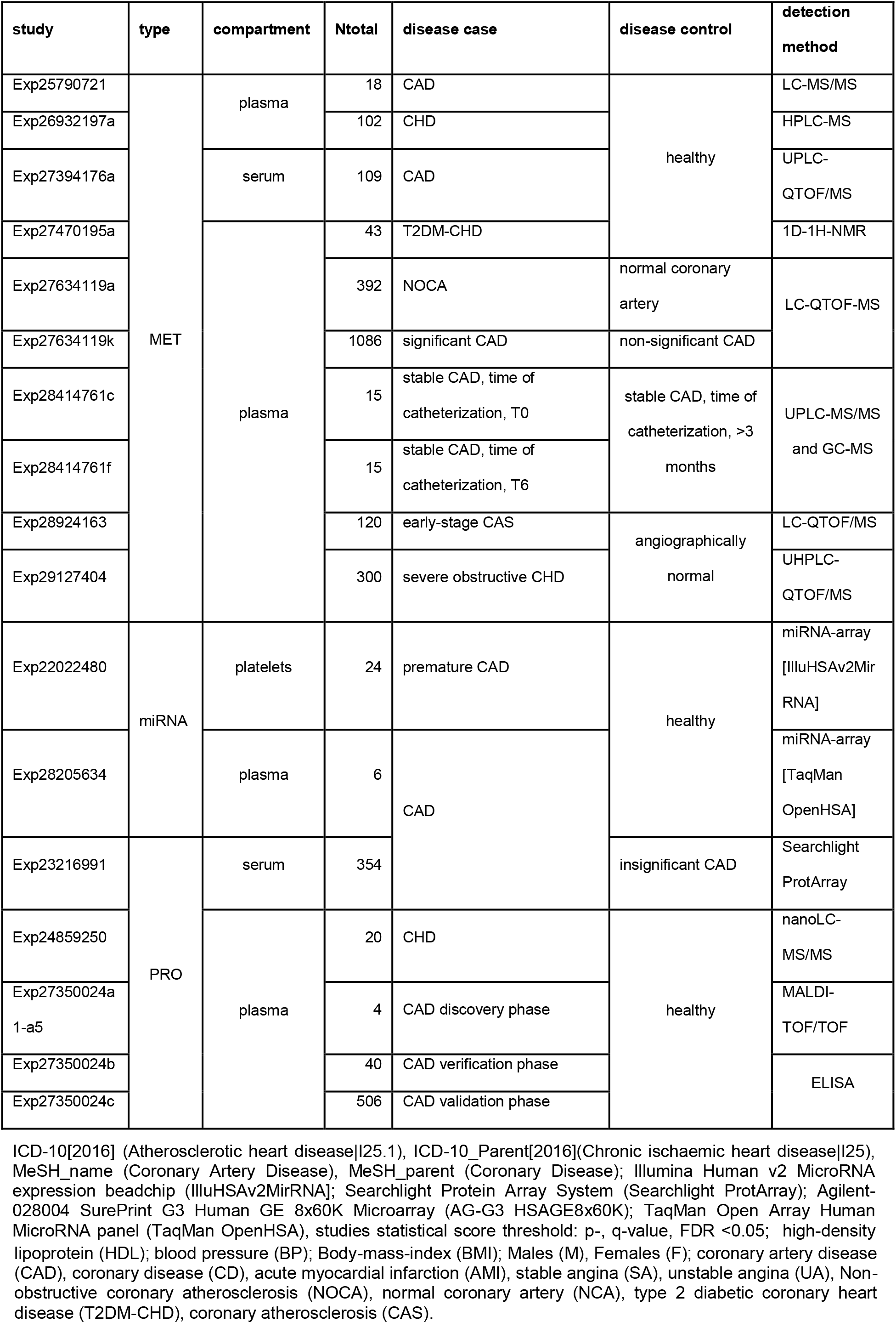
Dataspace description of CAD studies, from experimental design to clinical information. Human studies detected in blood.

ICD-10[2016] (Atherosclerotic heart disease|I25.1), ICD-10_Parent[2016](Chronic ischaemic heart disease|I25), MeSH_name (Coronary Artery Disease), MeSH_parent (Coronary Disease); Illumina Human v2 MicroRNA expression beadchip (IlluHSAv2MirRNA]; Searchlight Protein Array System (Searchlight ProtArray); Agilent-028004 SurePrint G3 Human GE 8×60K Microarray (AG-G3 HSAGE8×60K); TaqMan Open Array Human MicroRNA panel (TaqMan OpenHSA), studies statistical score threshold: p-, q-value, FDR <0.05; high-density lipoprotein (HDL); blood pressure (BP); Body-mass-index (BMI); Males (M), Females (F); coronary artery disease (CAD), coronary disease (CD), acute myocardial infarction (AMI), stable angina (SA), unstable angina (UA), Non-obstructive coronary atherosclerosis (NOCA), normal coronary artery (NCA), type 2 diabetic coronary heart disease (T2DM-CHD), coronary atherosclerosis (CAS).

The frequency (%) distribution of the regulation (down, up and not regulated) across three fold-change (FC) thresholds (|FC|>1.3, 1.5 and 2.0) of the most reported (at least two times) in CAD of microRNA (MIR), protein (PRO) and metabolite (MET) is shown in table 2. In any case of contradictory regulation, prevalence was given to any observing frequency ≥60%. The expression matrix handling differential expression raw values is available in supplementary table 2.

**Table 2.** Most reported molecular entities in CAD. Overall reported p-value <0.05 and frequency (%) of their differential expression/regulation (DOWN or UP) in each threshold group.

|  | Frequency (%) regulation |  |  |  |  |  |  |  |  | Reported | Tag |  |
| --- | --- | --- | --- | --- | --- | --- | --- | --- | --- | --- | --- | --- |
|  | \|FC\|>1.3 |  |  | \|FC\|>1.5 |  |  | \|FC\|>2.0 |  |  |  |  |  |
| Name | DOWN | UP | NR | DOWN | UP | NR | DOWN | UP | NR |  |  |  |
| hsa-mir-144 | 50 | 0 | 50 | 0 | 0 | 100 | 0 | 0 | 100 | 2 | MIR |  |
| hsa-miR-505-3p | 50 | 0 | 50 | 0 | 0 | 100 | 0 | 0 | 100 | 2 |  |  |
| hsa-let-7c | 0 | 50 | 50 | 0 | 50 | 50 | 0 | 50 | 50 | 2 |  |  |
| hsa-mir-1303 | 0 | 50 | 50 | 0 | 50 | 50 | 0 | 50 | 50 | 2 |  |  |
| hsa-mir-1305 | 0 | 50 | 50 | 0 | 50 | 50 | 0 | 50 | 50 | 2 |  |  |
| hsa-mir-943 | 0 | 50 | 50 | 0 | 50 | 50 | 0 | 50 | 50 | 2 |  |  |
| hsa-miR-579-3p | 0 | 50 | 50 | 0 | 50 | 50 | 0 | 50 | 50 | 2 |  |  |
| hsa-miR-10a-5p | 50 | 50 | 0 | 50 | 0 | 50 | 50 | 0 | 50 | 2 |  |  |
| hsa-mir-362 | 50 | 0 | 50 | 50 | 0 | 50 | 50 | 0 | 50 | 2 |  |  |
| hsa-mir-20a | 50 | 0 | 50 | 50 | 0 | 50 | 50 | 0 | 50 | 2 |  |  |
| hsa-mir-576 | 50 | 50 | 0 | 50 | 0 | 50 | 50 | 0 | 50 | 2 |  |  |
| hsa-miR-106b-5p | 100 | 0 | 0 | 50 | 0 | 50 | 50 | 0 | 50 | 2 |  |  |
| hsa-miR-517a-3p | 100 | 0 | 0 | 50 | 0 | 50 | 50 | 0 | 50 | 2 |  |  |
| hsa-mir-636 | 50 | 0 | 50 | 50 | 0 | 50 | 50 | 0 | 50 | 2 |  |  |
| hsa-mir-661 | 50 | 0 | 50 | 50 | 0 | 50 | 50 | 0 | 50 | 2 |  |  |
| hsa-mir-132 | 50 | 50 | 0 | 50 | 0 | 50 | 50 | 0 | 50 | 2 |  |  |
| hsa-miR-21-5p | 50 | 0 | 50 | 50 | 0 | 50 | 50 | 0 | 50 | 2 |  |  |
| hsa-miR-1255b-5p | 50 | 0 | 50 | 50 | 0 | 50 | 50 | 0 | 50 | 2 |  |  |
| APOB | 28.6 | 0 | 71 | 14.3 | 0 | 86 | 0 | 0 | 100 | 7 | TP | PRO |
| APOA4, ApoA-IV, APOA-IV | 66.7 | 0 | 33 | 16.7 | 0 | 83 | 16.7 | 0 | 83 | 6 | TP |  |
| TMSB4X, TB4X, THYB4 | 0 | 80 | 20 | 0 | 80 | 20 | 0 | 60 | 40 | 5 | CS |  |
| APCS, PTX2 | 80 | 0 | 20 | 60 | 0 | 40 | 40 | 0 | 60 | 5 | DIS |  |
| ADIPOQ, ACDC, ACRP30 | 60 | 0 | 40 | 0 | 0 | 100 | 0 | 0 | 100 | 5 | CS |  |
| APOA1 | 80 | 0 | 20 | 60 | 0 | 40 | 0 | 0 | 100 | 5 | TP |  |
| PRDX2, NKEFB, TDPX1 | 0 | 0 | 100 | 0 | 0 | 100 | 0 | 0 | 100 | 4 | ENZ |  |
| APOC2, APC2 | 33.3 | 33 | 33 | 33.3 | 33 | 33 | 0 | 0 | 100 | 3 | TP |  |
| VTN | 0 | 67 | 33 | 0 | 0 | 100 | 0 | 0 | 100 | 3 | CS |  |
| GPX3, GPXP | 33.3 | 0 | 67 | 33.3 | 0 | 67 | 33.3 | 0 | 67 | 3 | ENZ |  |
| APOC1, APOC1B | 33.3 | 0 | 67 | 33.3 | 0 | 67 | 33.3 | 0 | 67 | 3 | TP |  |
| ACTG1, ACTG | 0 | 100 | 0 | 0 | 100 | 0 | 0 | 50 | 50 | 2 | CS |  |
| ALB, GIG20, GIG42 | 0 | 0 | 100 | 0 | 0 | 100 | 0 | 0 | 100 | 2 | TP |  |
| AMBP, ITIL, HCP | 0 | 100 | 0 | 0 | 50 | 50 | 0 | 0 | 100 | 2 | INH |  |
| APOL1, APOL | 50 | 0 | 50 | 50 | 0 | 50 | 0 | 0 | 100 | 2 | TP |  |
| MPO | 50 | 50 | 0 | 50 | 0 | 50 | 50 | 0 | 50 | 2 | ENZ |  |
| LysoPC(18:2(9Z,12Z)) | 75 | 25 | 0 | 0 | 25 | 75 | 0 | 25 | 75 | 4 | MET |  |
| LysoPC(22:6(4Z,7Z,10Z,13Z,16Z,19Z)) | 50 | 25 | 25 | 25 | 25 | 50 | 0 | 25 | 75 | 4 |  |  |
| LysoPC(18:1(9Z)) | 75 | 25 | 0 | 25 | 25 | 50 | 0 | 25 | 75 | 4 |  |  |
| LysoPE(0:0/18:2(9Z,12Z)) | 75 | 25 | 0 | 0 | 25 | 75 | 0 | 25 | 75 | 4 |  |  |
| LysoPC(18:3(9Z,12Z,15Z)) | 75 | 0 | 25 | 25 | 0 | 75 | 0 | 0 | 100 | 4 |  |  |
| LysoPC(14:0/0:0) | 75 | 0 | 25 | 25 | 0 | 75 | 25 | 0 | 75 | 4 |  |  |
| LysoPC(18:0) | 75 | 0 | 25 | 50 | 0 | 50 | 25 | 0 | 75 | 4 |  |  |

|  |  |  |  |  |  |  |  |  |  |  |
| --- | --- | --- | --- | --- | --- | --- | --- | --- | --- | --- |
| LysoPC(16:1(9Z)/0:0) | 100 | 0 | 0 | 50 | 0 | 50 | 25 | 0 | 75 | 4 |
| LysoPE(16:0/0:0) | 50 | 0 | 50 | 0 | 0 | 100 | 0 | 0 | 100 | 4 |
| LysoPC(20:4(5Z,8Z,11Z,14Z)) | 25 | 0 | 75 | 0 | 0 | 100 | 0 | 0 | 100 | 4 |
| LysoPC(18:4(6Z,9Z,12Z,15Z)) | 75 | 25 | 0 | 0 | 25 | 75 | 0 | 25 | 75 | 4 |
| LysoPC(15:0) | 66.7 | 33 | 0 | 33.3 | 33 | 33 | 0 | 33 | 67 | 3 |
| Phytosphingosine | 66.7 | 33 | 0 | 66.7 | 0 | 33 | 33.3 | 0 | 67 | 3 |
| Palmitic acid | 66.7 | 33 | 0 | 33.3 | 0 | 67 | 33.3 | 0 | 67 | 3 |
| LysoPE(0:0/18:0) | 33.3 | 0 | 67 | 0 | 0 | 100 | 0 | 0 | 100 | 3 |
| LysoPE(20:4(5Z,8Z,11Z,14Z)/0:0) | 0 | 33 | 67 | 0 | 0 | 100 | 0 | 0 | 100 | 3 |
| L-tyrosine | 50 | 50 | 0 | 0 | 0 | 100 | 0 | 0 | 100 | 2 |
| Dehydroepiandrosterone sulfate (DHEA) | 50 | 50 | 0 | 0 | 50 | 50 | 0 | 0 | 100 | 2 |
| L-arginine | 50 | 50 | 0 | 0 | 50 | 50 | 0 | 50 | 50 | 2 |
| LysoPE(18:2(9Z,12Z)/0:0) | 50 | 50 | 0 | 0 | 50 | 50 | 0 | 50 | 50 | 2 |
| Alpha-linolenic acid (ALA) | 50 | 50 | 0 | 50 | 0 | 50 | 50 | 0 | 50 | 2 |
| N-Palmitoylethanolamide (PEA) | 100 | 0 | 0 | 100 | 0 | 0 | 0 | 0 | 100 | 2 |
| Cer(d18:0/16:0) | 0 | 100 | 0 | 0 | 50 | 50 | 0 | 50 | 50 | 2 |
| Pregnenolone sulfate | 100 | 0 | 0 | 100 | 0 | 0 | 50 | 0 | 50 | 2 |
| PC(16:0/0:0)[rac] | 0 | 0 | 100 | 0 | 0 | 100 | 0 | 0 | 100 | 2 |
| PE(21:0/0:0) | 0 | 0 | 100 | 0 | 0 | 100 | 0 | 0 | 100 | 2 |
| LysoPC(22:5(7Z,10Z,13Z,16Z,19Z)) | 50 | 0 | 50 | 0 | 0 | 100 | 0 | 0 | 100 | 2 |
| L-valine | 0 | 50 | 50 | 0 | 0 | 100 | 0 | 0 | 100 | 2 |
| LysoPE(18:1(9Z)/0:0) | 100 | 0 | 0 | 0 | 0 | 100 | 0 | 0 | 100 | 2 |
| LysoPE(18:0/0:0) | 0 | 0 | 100 | 0 | 0 | 100 | 0 | 0 | 100 | 2 |
| LysoPE(0:0/20:1(11Z)) | 50 | 0 | 50 | 0 | 0 | 100 | 0 | 0 | 100 | 2 |
| 2-Hydroxybutyric acid | 0 | 50 | 50 | 0 | 0 | 100 | 0 | 0 | 100 | 2 |
| L-Acetylcarnitine (ALCAR or ALC) | 0 | 50 | 50 | 0 | 0 | 100 | 0 | 0 | 100 | 2 |
| Glycine | 50 | 0 | 50 | 0 | 0 | 100 | 0 | 0 | 100 | 2 |
| LysoPE(0:0/22:5(4Z,7Z,10Z,13Z,16Z)) | 50 | 0 | 50 | 0 | 0 | 100 | 0 | 0 | 100 | 2 |
| L-histidine | 50 | 0 | 50 | 0 | 0 | 100 | 0 | 0 | 100 | 2 |
| LysoPC(22:5(4Z,7Z,10Z,13Z,16Z)) | 50 | 50 | 0 | 0 | 0 | 100 | 0 | 0 | 100 | 2 |
| Cortisol | 0 | 50 | 50 | 0 | 50 | 50 | 0 | 50 | 50 | 2 |
| Docosahexaenoic acid (DHA) | 50 | 0 | 50 | 50 | 0 | 50 | 50 | 0 | 50 | 2 |
| 8,11,14-Eicosatrienoic acid (DGLA) | 50 | 0 | 50 | 50 | 0 | 50 | 50 | 0 | 50 | 2 |
| Icosanoic acid (eicosanoic acid) | 50 | 50 | 0 | 50 | 0 | 50 | 50 | 0 | 50 | 2 |
| 1,7-dimethylxanthine (paraxanthine) | 50 | 50 | 0 | 50 | 0 | 50 | 50 | 0 | 50 | 2 |

Not regulated (NR). Fold-change (FC). MIR: microRNA, TP: transport, storage, endocytosis, exocytosis, vesicles, CS: Cell shape (cytoskeleton, cell adhesion, morphology, cell junction, cellular structures, extracellular matrix), DIS: disease, ENZ: enzyme, enzymatic properties, INH: inhibitor (protease, kinase, other enzymes, pathways), and MET: metabolite.

The 76 most reported molecules in CAD which includes 18 microRNAs, 16 proteins and 42 metabolites (Table 2) were analysed either by gene ontology (GO), for their biological process (BP), molecular function (MF) and cellular component (CC) and/or Kyoto encyclopedia of genes and genomes (KEGG-compounds) (Fig 5).

**Fig 5.**
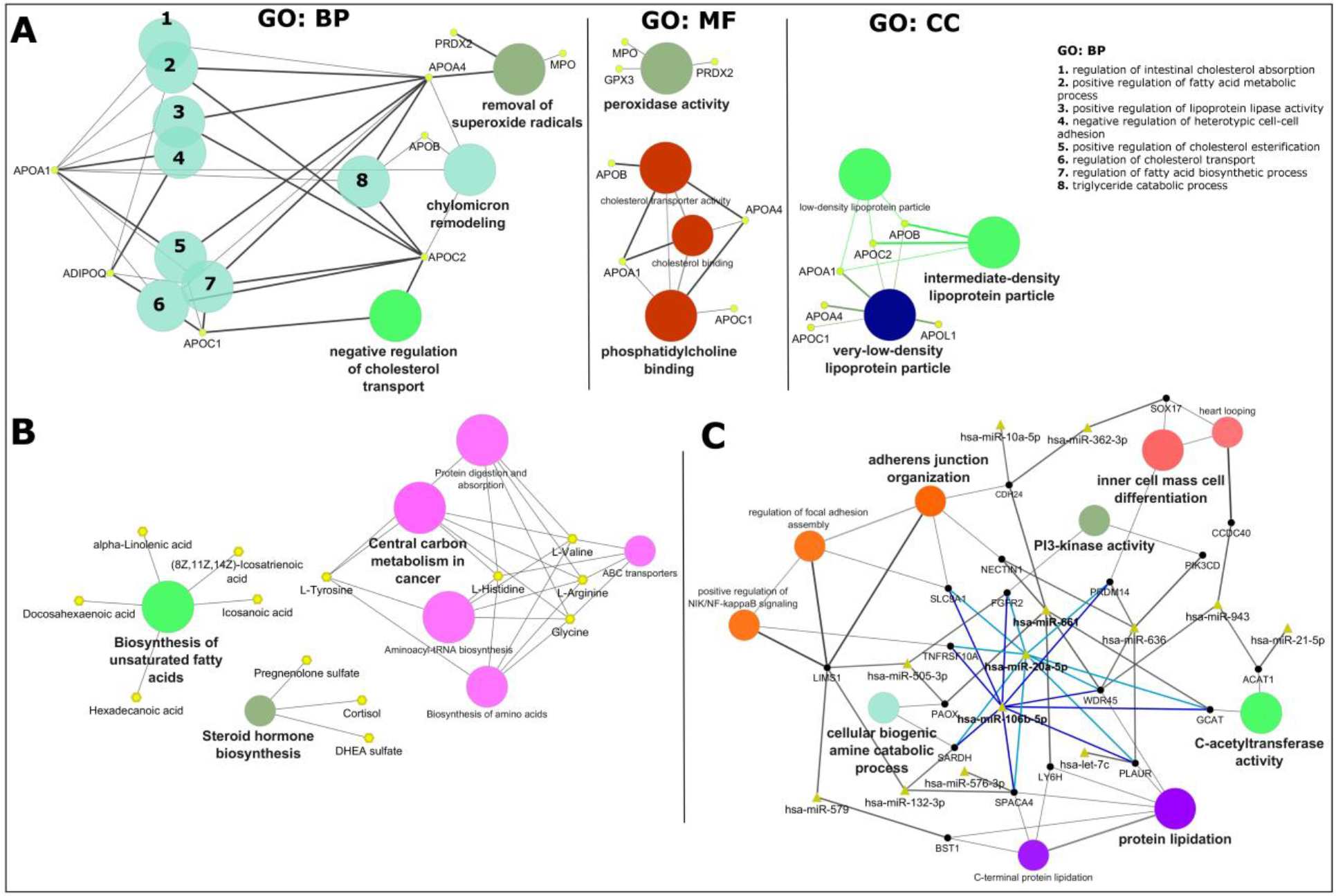
Exploratory analysis of CAD using the most reported molecular entities. A: Gene ontology (GO) analysis of the proteomics data sets. Biological process (BP), molecular function (MF) and cell component (CC). Node size of GO/pathway terms corresponds to the number of matched genes/proteins. B: compound analysis using KEGG pathway maps. C: miRNA analysis of the respective gene target(s) using miRanda predictions. miRNA-targets were then converged onto GO terms. In A: apolipoprotein A-I (APOA1), adiponectin (ADIPOQ), apolipoprotein C-I (APOC1), apolipoprotein B-100 (APOB), apolipoprotein A-IV (APOA4), myeloperoxidase (MPO), peroxiredoxin-2 (PRDX2), glutathione peroxidase 3 (GPX3), apolipoprotein C-II (APOC2), and apolipoprotein L1 (APOL1). In B: hexadecanoic acid (palmitic acid), (8Z,11Z,14Z)-icosatrienoic acid, alpha-linolenic acid (ALA), docosahexaenoic acid (DHA), icosanoic acid (eicosanoic acid), DHEA sulfate (dehydroepiandrosterone sulfate), pregnenolone sulfate (3beta-Hydroxypregn-5-en-20-one sulfate). In C: phosphatidylinositol 4,5-bisphosphate 3-kinase catalytic subunit delta isoform (PIK3CD), acetyl-CoA acetyltransferase, mitochondrial WD repeat domain phosphoinositide-interacting protein 4 (WDR45), sperm acrosome membrane-associated protein 4 (SPACA4), sarcosine dehydrogenase, mitochondrial (SARDH), peroxisomal N(1)-acetyl-spermine/spermidine oxidase (PAOX), fibroblast growth factor receptor 2 (FGFR2), 2-amino-3-ketobutyrate coenzyme A ligase, mitochondrial (GCAT), acetyl-CoA acetyltransferase, mitochondrial (ACAT1), LIM and senescent cell antigen-like-containing domain protein 1 (LIMS1), lymphocyte antigen 6H (LY6H), phosphatidylinositol 4,5-bisphosphate 3-kinase catalytic subunit delta isoform (PIK3CD), urokinase plasminogen activator surface receptor (PLAUR), coiled-coil domain-containing protein 40 (CCDC40), nectin-1 (NECTIN1), PR domain zinc finger protein 14 (PRDM14), transcription factor SOX-17 (SOX17), cadherin-24 (CDH24), sodium/hydrogen exchanger 1 (SLC9A1), ADP-ribosyl cyclase/cyclic ADP-ribose hydrolase 2 (BST1), and tumor necrosis factor receptor superfamily member 10A (TNFRSF10A).

GO cluster analysis using ClueGO (Fig 5A) showed that the most reported proteins in CAD are primarily involved in the biological regulation of cholesterol transport (negative regulation (APOC1 and APOC2)) and absorption (intestinal absorption (APOA1 and APO4)), modification of the chylomicron composition (APOB, APOC2, APOA1 and APOA4), regulation of metabolic processes regarding fatty acid biosynthesis (APOC1, positive regulation (ADIPOQ, APOA1, APOA4 and APOC2)) and cellular detoxification of superoxide radicals via peroxidase activity (APOA4, MPO, GPX3 and PRDX2). These molecules consist mostly of plasma lipoprotein particles (very-low-density (APOL1, APOC1, APOA4, APOA1, APOC2 and APOB), low-density (APOA1, APOC2 and APOB) and intermediate-density (APOB, APOC2 and APOA1)). The nine characterized proteins clustered in the network follow a trend of decreased expression so we can expect the same trend for their associated functions and biological processes described above. Additionally, ClueGO analysis of proteins reported once (Supplemental Fig S6A), and their connectivity based on STRING protein-protein interactions (Supplemental Fig S6B), uncovered the PPAR signalling pathway (containing ADIPOQ, FABP1 and APOA1), extracellular matrix (ECM)-receptor interaction (ITGB3, SPP1, CD44 and VTN) activation of the complement and proteolytic coagulation cascades (FGA, FGG, KNG1 and SERPIND1) clusters and highlighted the importance of the statins pathway with the cluster (APOC2, LCAT, APOB, APOA4, APOC, APOA1, ADIPOQ and FABP1).

Similarly, compound analysis using the Kyoto encyclopedia of genes and genomes (KEGG-compounds) in ClueGO (Fig 5B) yielded the processes of biosynthesis of unsaturated fatty acids (hexadecanoic acid, (8Z,11Z,14Z)-icosatrienoic acid, alpha-linolenic acid, docosahexaenoic acid and icosanoic acid), steroid hormone biosynthesis (pregnenolone sulfate, cortisol and DHEA sulfate) and central carbon metabolism (L-tyrosine, L-histidine, L-valine, L-arginine and glycine) with child elements as biosynthesis of amino acids, ABC transporters, aminoacyl-tRNA biosynthesis and involved in protein digestion and absorption. Most of the metabolites, nine out of thirteen were found decreased in expression (exception for L-valine, L-arginine, cortisol and DHEA sulfate) so one can expect the same trend for their associated biological and metabolic processes.

Following the rationale that microRNAs exert post-transcriptional regulation upon mRNA/gene targets, thereby inversely regulated miR-target pairs can contribute to either an increased or decreased expression of the target. Therefore, based on the network cluster (Fig 5C) that connects microRNAs-targets via miRanda target prediction (using the CluePedia 1.5.0 app) to biological processes and molecular functions and observing the regulation trend of microRNAs in table 2, we can verify that 10 out of 13 microRNAs displayed in the network are decreased in expression (exception for miR-579, let-7c and miR-943), thus one can expect that their targets would be increased in expression, so as their associated biological process. Specifically, as seen with miR-661, miR-20a-5p and miR-106b-5p that have a greater number of targets and follow a down-regulation trend, their targets are involved in protein lipidation, C-acetyltransferase activity, adherens junction organization and cellular biogenic amine catabolic process for the first described microRNA and additionally the PI3-kinase activity and inner cell mass cell differentiation for the two latter described microRNAs (Fig 5C). Additional, search of potential affected KEGG pathways in mirPath v3.0 yield the fatty acid biosynthesis pathway (KEGG map, hsa00061), with up-regulated microRNAs, miR-1303 and miR-1305 targeting FASN.

#### Interactome analysis and network enrichment

We established a global interactome network (Fig 6) starting with the most reported proteins in CAD, making associations based on protein-protein interactions using STRING app. Afterwards, as a gap-filling approach, the least reported proteins (supplementary Excel file) were added, as well as the highest scored gene associations from the DisGeNET database (supplementary Excel file). Furthermore, we incorporated metabolic associations providing the linkage of gene-enzyme-reaction-metabolite by merging the former network with a metabolic network built in MetScape (Supplemental Fig S7), and described by ClueGO analysis (Supplemental Fig S8 and S9). Later, we kept only the most reported and regulated microRNAs (miRNAs), providing their association with gene targets present in the network via CluePedia. The possible occurrence of other regulatory elements in the recently developed network such as transcription factors (TF’s) was ensured by the use of CyTargetLinker and GeneMANIA.

**Fig 6.**
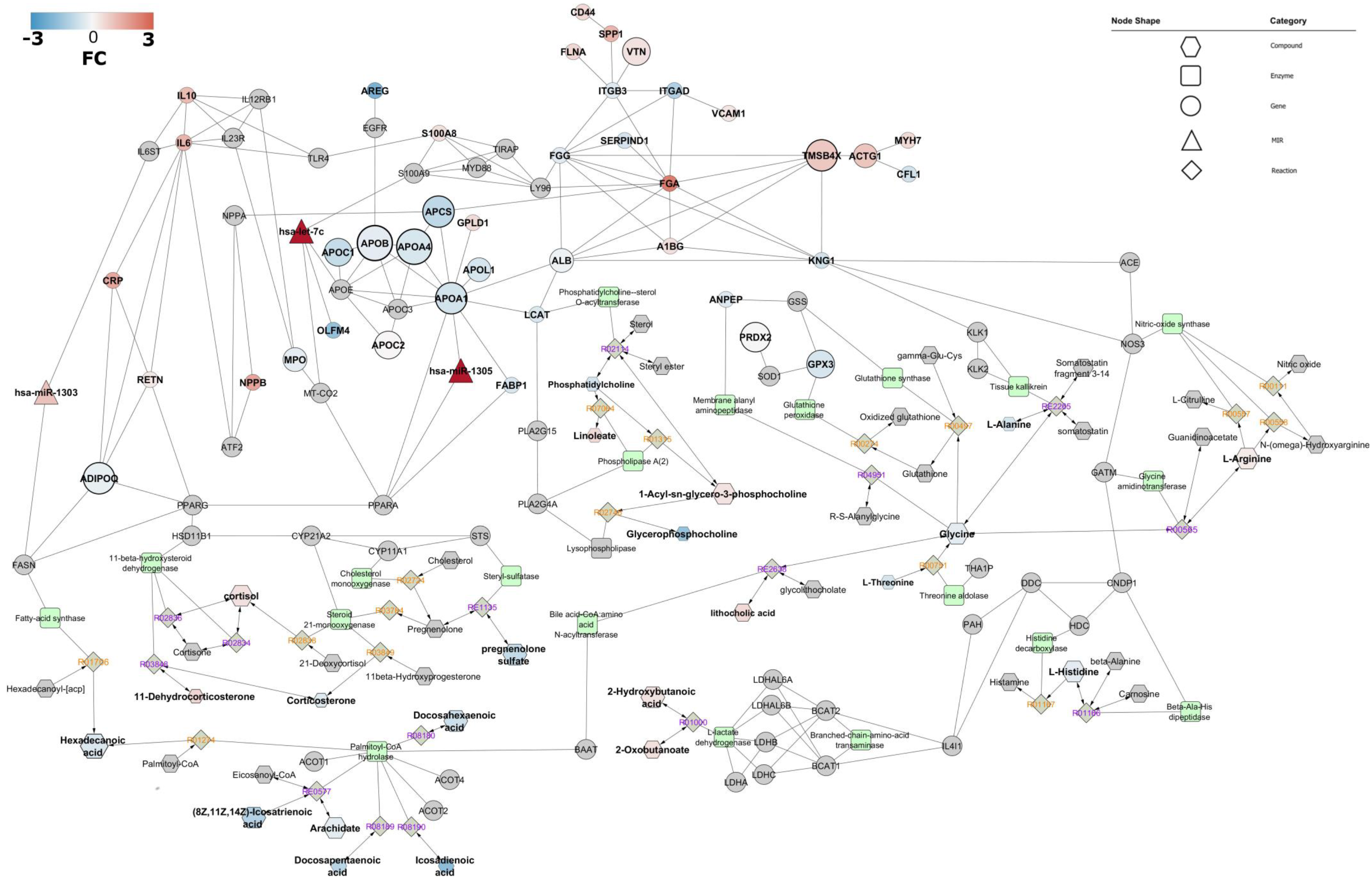
CAD interactome. Network representation of microRNAs as node triangles, proteins/genes as circles, enzymes displayed as round rectangles, compounds as hexagons, and reactions shaped as diamonds, labels colour coded as irreversible/directed: orange, reversible/bidirected: purple. Enrichment using protein-protein physical interactions is derived from STRING with an established minimum confidence score of 0.70. Gene-enzyme-reaction-compound associations were established using MetScape 3.1.3 and KEGG. MicroRNA-targets associations were derived using miRanda and TarBase. Differential expression is represented as a colour gradation from blue (decreased expression) to red (increased expression). Node size is proportional to the number of molecules reported within data sets. Some compounds and microRNAs of the network were removed (based on fold-change criteria) to reduce the complexity of the figure and thereby enhance visualisation. Full network figure is available in Supplementary Fig S10.

This resulted in a densely interconnected network (Fig 6). This network of 136 entries (expanded full network contains 176 molecular entities, Supplementary Fig S10), including molecules known to be associated with the query-molecules, consists of a sub-expressed apolipoprotein-cluster (containing APOC1, APOB, APOA4, APOA1, APOC2 and APOL1) and LCAT, a main driver of the extracellular metabolism of lipoproteins, involvement of the complement and coagulation cascades (FGA, FGG, SERPIND1, KNG1 and VTN), suggesting inflammation being activated in the original source tissue, and processes including extracellular matrix receptor interaction (ITGB3, SPP1, CD44 and VTN), biosynthesis of unsaturated fatty acids (hexadecanoic acid, (8Z,11Z,14Z)-icosatrienoic acid, alpha-linolenic acid, docosahexaenoic acid, docosapentaenoic acid, icosanoic acid, arachidate and oleate), steroid hormone biosynthesis (pregnenolone sulfate, cortisol, corticosterone, 11-dehydrocorticosterone and DHEA sulfate), amino acid metabolism (L-arginine, L-histidine, glycine and L-alanine) transport (ALB, APCS), cellular detoxification of oxygen species (PRDX2, MPO and GPX3) and actin cytoskeleton (CFL1, ITGAD, ITGB3, ACTG1, TMSB4X). Inter-linking molecules suggest an involvement of transcriptional elements such as ATF2, PPARG, PPARA and EGFR, which are modulators of genes involved in DNA damage, cell proliferation, anti-apoptosis, glucose and lipid metabolic processes. Additionally, keeping only microRNAs highly expressed and inversely correlated with targets, yielded regulatory clusters driven by miR-1305 (directly targeting APOA1 and PPARA), miR-let-7c (directly targeting OLFM4, and resulting query-molecules S100A9 and MT-COX2/COX2) and miR-1303 (targeting both resulting query-molecules IL6ST and FASN).

Analysis of the condensed network of direct interactions between the 136 (or 176 proteins, full network, Supplementary Fig S10) proteins shows the apolipoprotein-cluster containing APOC1, APOB, APOA4, APOA1, APOC2 and APOL1, also associated with ALB and LCAT, more than three interactions between IL6, Il10, RETN, CRP, and ADIPOQ and also between FABP1 and the apolipoprotein-cluster hub APOA1. The latter’s suggests that the peroxisome proliferator-activated receptor (PPAR) signalling pathway is perturbed in CAD through an association with APOA1, FABP1 (PPARA), ADIPOQ and RETN (PPARG) (Figure 6). Additional, manual queries in the C/VD database (human, CAD or atherosclerosis, detected in blood, gene expression studies, FC>1.5) looking for regulation trends of network molecules with absent regulation, resulted in decreased levels of PPARG, MT-CO2/COX2, and FASN, confirming the general trend of the PPAR signalling pathway to be inactivated/supressed/blunted.

Data mining of the DisGeNET database by associated disease clustering having as input the 16 proteins from table 2 showed association of MPO, APOA1, APOB and ADIPOQ with myeloperoxidase deficiency, hypertensive disease, hypercholesterolemia, hyperlipidemia and hypoalphalipoproteinemia. In a similar approach to mine pathway entries in the KEGG database, the common denominator was found as hypertrophic cardiomyopathy (HCM, KEGG entry hsa05410) containing the cluster IL6, ITGB3 and ACTG1 (table 3), linking it to the ECM-receptor interaction, renin-angiotensin system JASK-STAT signalling pathway and TGFbeta-signalling pathway.

**Table 3.**
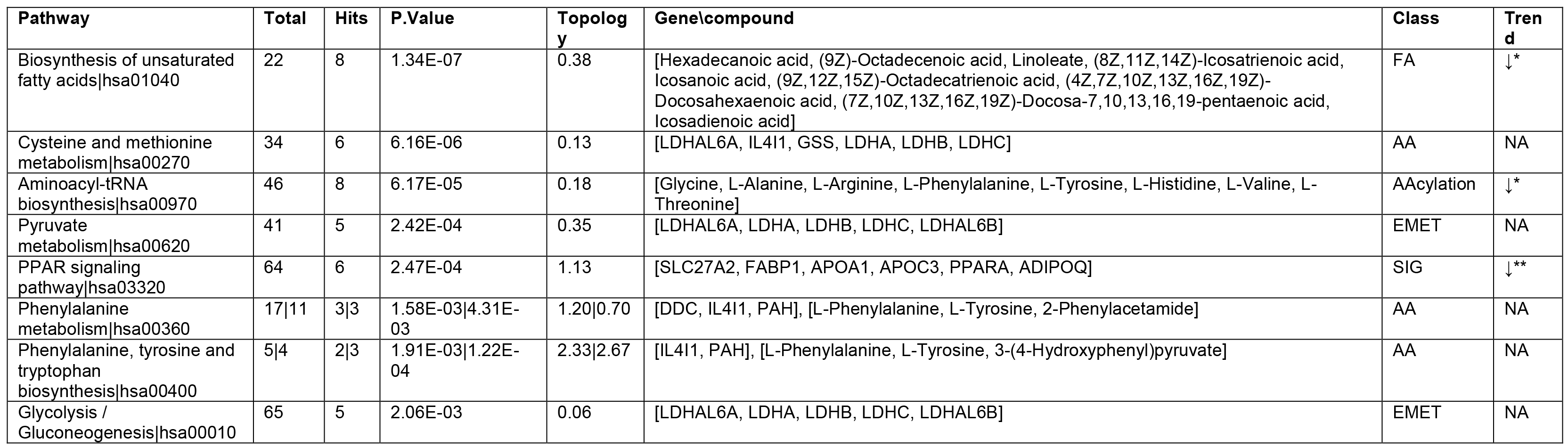
Over-representation and pathway topology analysis using KEGG pathways terms. The analysis is based on the molecular elements that constitute the interactome of CAD. Topology analysis is based on degree centrality, which measures the number of links that connects to a node within a pathway. Regulation trend (Trend) is based on differential expression of either *compounds, or **gene\proteins, or ***both.

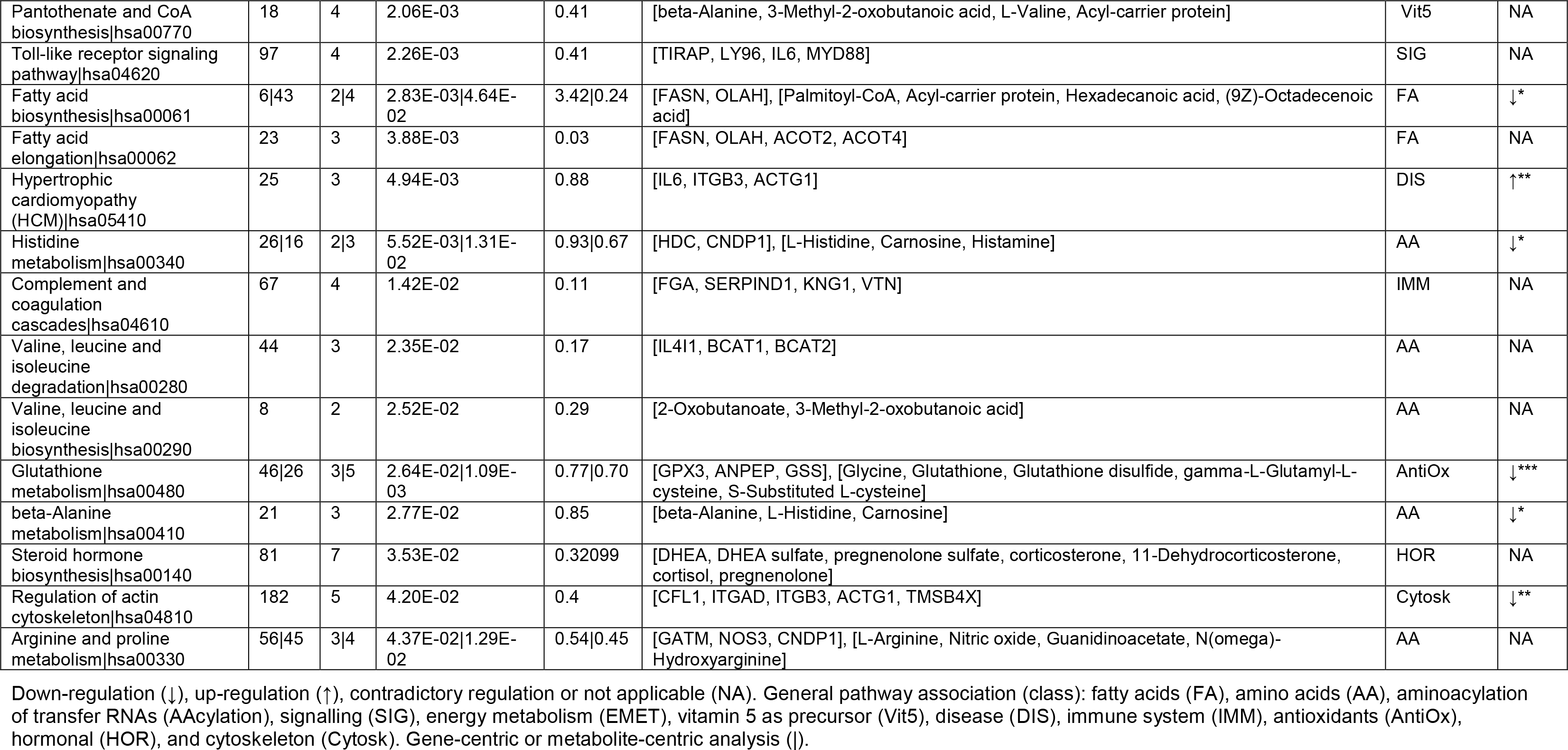

## Discussion

The C/VD database handles multi-omics and clinical data retrieved from published experimental studies in literature and by mining NCBI Gene Expression Omnibus (GEO) [29] resources across multi-species and tissue/fluid sources in cardiovascular disease. Systems-level integration requires not only large-scale data, but mostly comprehensive data at all molecular levels, thereby owning a resource able to summon all of these elements altogether on a simple, comparative and clustered platform is of prime importance. Moreover, as C/VD is part of the Pan-omics Analysis Database (PADB, www.padb.org) framework, its stability over-time and regular update is ensured, which is a real problem that many databases face after publication.

Databases covering cardiovascular disease molecular data exist, in the vast majority only covering single-omics derived data, with prevalence for gene-array data sourced from GEO, furthermore many are either offline or their domain redirects to other unrelated webpages. CardioGenBase [61] displays gene-array data sourced from literature and GEO and its currently offline (verified on 05/07/2018), another database comprising literature-based candidate CAD genes, CADgene, [62] and its database live status is currently offline (verified on 05/07/2018, redirects to another unrelated web-page), likewise, In-Cardiome [63] deals with own-private and literature-based gene expression data, as well with clinical associations and drugs; Chemogenomics [64] is a literature-based database aiming to explore more extensively the pharmacological side in cardiovascular disease handling protein targets and small ligands, and is currently online (verified on 05/07/2018), but requires registration; COPaKB [65] mimics The Human Protein Atlas database [66] with a subset of data regarding cardiovascular disease and also GEO data, and its live status is online (verified on 05/07/2018).

The generated interactome in coronary disease (CAD) based on initial meta-analysis and further systems-level integrative analysis of Omics studies covering circulatory levels of protein, metabolite and microRNA differential expression spotted disturbed profiles in lipid metabolism, in particular down-regulation of the unsaturated fatty acids (FA) biosynthesis, down-regulation of cholesterol binding and transport ability shown by the apolipoprotein containing cluster, phosphatidylcholine-sterol acyltransferase (LCAT) and (FABP1), the involvement of the proliferator-activated receptor (PPAR) signalling pathway, with particular emphasis on the generalised trend for down-regulation of the peroxisome proliferator-activated receptor gamma (PPARG), fatty acid synthase (FASN) and their enzyme product, hexadecanoic acid (palmitic acid). This is in synergy with putative post-transcriptional regulation of miR-1305 on PPARA and APOA1, a main apolipoprotein cluster network hub, and dual regulation of miR-1303 and miR-1305 of FASN. On the other hand, indication of up-regulation of molecular elements involved in the extracellular matrix (ECM)-receptor interaction, inflammation being activated in the original source tissue and cardiac hypertrophy.

Peroxisome proliferator-activated receptors (PPARs) alpha, beta/delta and gamma are ligand activated transcription factors that share a substantial homology, 60% to 80%, in their ligand and DNA-binding domains [67], despite playing different roles in the modulation of energy metabolism [68]. The three PPARs isoforms are expressed in the heart, including vasculature of endothelial [69, 70] and smooth muscle cells [71, 72], but their roles in the cardiovascular system and outcomes in disease are still not well established due to major inter-study variability [73]. The activation or repression of the expression of PPARs gene targets is modulated by transactivation or transrepression mechanisms [74], in which binding of ligands to PPAR leads to the establishment of heterodimers with the nuclear retinoid receptor (RXR), followed by translocation of PPAR-RXR-heterodimer-complex to the nucleus, with subsequent assembling of co-activator (e.g. PPARGC1A) complexes [75] and binding to specific PPAR response elements (PPREs) in the promotor region of target genes to induce their expression [70, 76]. In contrast, in the absence of ligands, the PPAR-RXR-heterodimer-complex recruits co-repressors (e.g. NCOR1 and NCOR2) that repress gene expression in a DNA-binding-independent manner. When a ligand is present and binds, PPAR-RXR induces dissociation of the co-repressors complex and subsequent release of the co-repressor elements [70, 76, 77].

PPARs are implicated in a plethora of cardiovascular disorders such as cardiac hypertrophy, atherosclerosis and heart failure, and as well in conditions that pose a risk factor, such as diabetes mellitus, obesity, hypertension and dyslipidemia [78].

Activation of PPARA by binding of fibric acid derivatives (fibrates) results in augmented levels of high-density lipoprotein (HDL) cholesterol, diminished levels of free fatty acids (FFA), triglycerides, apolipoprotein C-III (APOC3) [71], and reduced inflammatory response via inhibition of NF-kappaB and transcription factor AP-1 (AP1/JUN) [79, 80]. In the liver, PPARA is involved in the degradation of fatty acids (FA) through the induction of numerous genes coding to enzymes participating in mitochondrial and peroxisomal fatty acid (FA) oxidation [81]. Furthermore, PPARA regulates gluconeogenesis [82], biotransformation [83] and cholesterol metabolism processes [83]. Natural occurring PPARA ligands comprise prostaglandins, leukotrienes, eicosapentaenoic acid and docosahexaenoic acid [84]. Many synthetic PPARA ligands were developed for the management of dyslipidaemia, covering several fibrate drug agonists as gemfibrozil, clofibrate, fenofibrate and bezafibrate [85].

PPARG activation often leads to processes associated with augmented insulin sensitivity, and perturbance of pancreatic beta-cell function [86], in adipocyte tissue PPARG enables fatty acid (FA) storage by inducing adipocyte differentiation [87], and up-regulates expression of the solute carrier family 2, facilitated glucose transporter member 4 (GLUT4/ SLC2A4) and phosphatidyl-3-kinase (PI3K) [88, 89]. Moreover, PPARG gene targets are associated with adipocyte differentiation, lipid storage and glucose metabolism, including lipoprotein lipase (LPL), platelet glycoprotein 4 (CD36), cytosolic phosphoenolpyruvate carboxykinase (PCK1), aquaporin-7 (AQP7) and adiponectin (ADIPOQ) [90]. Many unsaturated fatty acids bind to PPARG, including linoleic acid (linoleate) and arachidonic acid [89], and several eicosanoids [82]. PPARG is a known regulator of glucose metabolism, thereby its synthetic ligands includes many members of the antidiabetic thiazolidinedione (TZD) drug class, glitazones and telmisartan, an antagonist of the type-1 angiotensin II receptor (AGTR1/AT1) used in the treatment of hypertension [91]. Additionally, monocytes and macrophages can produce PPARG ligands, capable to induce an anti-inflammatory response and potentiate inflammatory pro-resolving actions towards repair and return to homeostasis [92].

Post-transcriptional regulators such as hsa-miR-1303, shown up-regulated profile when human umbilical vein endothelial cells (HUVECs) were exposed to estradiol [93], also a down-regulation signature of this microRNA when human-induced pluripotent stem cell-derived cardiomyocytes were exposed to doxorubicin [94]. Furthermore, hsa-miR-1305 levels in circulating monocytes was found to be increased after acute exercise as a stress signal in healthy volunteers [95], and hsa-let-7c associated with cardiac hypertrophy [96], and the mature sequence hsa-let-7c-5p appears to be associated with hyperglycemia in patients with coronary heart disease (CHD) [97].

Most patients with CAD undertake treatment for clinical management of other CAD risk factors, such as control of low-density lipoprotein (LDL) cholesterol levels, triglycerides and blood pressure, and therefore it is likely that the additional treatment can contribute to the global interactome profile highlighted above. Thus, we should empathise the potential role of statins (3-hydroxy-methylglutaryl coenzyme A, HMG-CoA reductase inhibitors), a group of drugs that are widely applied to reduce the level of blood low-density lipoprotein (LDL) cholesterol and triglycerides [98], and also found to increase the expression of PPARGC1A [99], a co-activator of PPARG. Other agonists as fibrates drugs are able to induce a diminished activity of LCAT in plasma [100].

The CAD showcase scenario presented here to demonstrate the utility of the C/VD database has some limitations as it only used data sets screening circulatory molecules in human cohorts, thereby potential disease mechanisms, disclosed in experimental studies using cell-lines and animal models mimicking CAD and as well processes mediated in specific tissue-types could not be fully perceived. Moreover, we selected only data sets dealing with the clinical description of CAD, leaving out early events perceived in conditions such as atherosclerosis and late diseases such as myocardial infarction, which for a molecular perspective can pose an incomplete view over all the involved molecular events undergoing this disease, thus leading to a partial description of CAD.

## Conclusions and future perspectives

The C/VD database can assist either on the development of novel hypotheses in a data-driven manner or be a source of literature-based knowledge in the cardiovascular research field.

In the CAD showcase, using a combinatorial systems-biology approach based on integration of data from circulatory molecular expression profiles covering microRNAs, protein and metabolite, we provide insights into the biological aspects, including summary descriptions by gene ontology (GO) and pathway terminology, covering mechanistic aspects through recreation of protein-protein interactions (PPI’s), metabolic reactions, and providing regulatory elements exerted by transcriptional factors (TF’s) and microRNAs post-transcriptional regulation. In CAD a global disturbed profile in lipid metabolism, including biosynthesis of fatty acids (FA), with blunted ability for cholesterol binding and transport, as well as the involvement of the proliferator-activated receptor (PPAR) signalling pathway, with PPAR isoform alpha (PPARA) being potentially post-transcriptionally regulated by miR-1305 and resulting sub-expression of PPARA target genes, APOA1 and FABP1. Furthermore, indication of disrupted biological processes with distinct increase in expression of molecular elements involved in activation of inflammation, extracellular matrix (ECM)-receptor interaction, and cardiac hypertrophy could be shown.

We foresee regular database updates, populating the C/VD database with more experimental studies since the cardiovascular research field, particularly large-scale untargeted approaches, screening body-accessible fluids are currently in high demand.

## Supporting information

Appendix file containing the following information:

**S1 Fig. Index page of the tissue/source list of the C/VD database.** The fields of disease, species, number of studies and number of molecules per each tissue/fluid (blood, heart, aorta, urine) source are described.

**S2 Fig. Index page for the omic studies detected in blood tissue/source.** The fields of species, component, number of molecules, disease, and PMID/DOI are described per study.

**S3 Fig. Structure of the C/VD database at the study page-view.** First headers enclose the main bibliometric data associated with a research article. The second and third headers are with respect to the description of the sample itself and several steps in their processing till reach molecule identification. The fourth header corresponds to the incoming molecules from the previous step. It lists all the molecules identified in the study with the associated statistical scores, fold-change and regulation.

**S4 Fig. View of the of a specific molecule entry in C/VD database.** In the first header is displayed the linkage to external databases. Following it we have represented the associated function and classification and as well our own type of tag for functional classification (derived from PADB). The third header corresponds to the number of occurrences of the molecule across all the studies, disease conditions, species, and tissue/fluid source.

**S5 Fig. Demographics page-view.** Clinical data (demographics page view) is also represented in the C/VD database and is linked with the study page and vice-versa.

**S6 Fig. Exploratory analysis of CAD using all reported proteins from the many proteomics data sets.** A: Gene ontology (GO) analysis. B: STRING protein-protein interactions with a minimum confidence score of 0.70. The light blue demarked oval in B denotes proteins from the hypertrophic cardiomyopathy (HCM) and ECM-receptor interactions ClueGO derived cluster. WikiPathways (WP), biological process (BP), molecular function (MF) and cell component (CC).

**S7 Fig. Visualisation of gene-enzyme-reaction-compound associations through MetScape.** Matched compounds and gene nodes are pink colour filled when derived from the C/VD database and grey when from DisGeNET regarding coronary artery disease (CAD) association.

**S8 Fig. Composite image of a MetScape enriched network (main cluster) followed by pathway association in ClueGO (KEGG compounds and pathway terms).** Most significant enriched node terms are displayed darker. Colour filled compounds and proteins/genes are from CAD datasets (exception for NOS3 that is derived from DisGeNET). To improve network visualisation some ontology terms were removed from the figure, such as protein digestion and absorption: blue stars; biosynthesis of amino acids: orange stars; central carbon metabolism in cancer: green stars.

**S9 Fig. Composite figure representing MetScape enriched network (remaining clusters) followed by pathway association in ClueGO (KEGG compounds and GO/pathway terms).** Most significant enriched node terms are displayed darker. Colour filled compounds and proteins/genes are from CAD datasets (exception for ACE that is derived from DisGeNET). To improve network visualisation some ontology terms were removed from the figure, such as pyruvate metabolism, glucagon signaling pathway, and glycolysis/gluconeogenesis that have a grey start annotation.

**S10 Fig. Full network representation of coronary artery disease (CAD) interactome.** Network representation of microRNAs as node triangles, proteins/genes as circles, enzymes displayed as round rectangles, compounds as hexagons, and reactions shaped as diamonds, labels colour coded as irreversible/directed: orange, reversible/bidirected: purple. Enrichment using protein-protein physical interactions is derived from STRING with an established minimum confidence score of 0.70. Gene-enzyme-reaction-compound associations were established using MetScape 3.1.3 and KEGG. MicroRNA-targets associations were derived using miRanda and TarBase. Differential expression is represented as a colour gradation from blue (decreased expression) to red (increased expression). Node size is proportional to the number of molecules reported within data sets.

**S1 Table. Clinical parameters of the human cohorts used in the CAD studies.** high-density lipoprotein (HDL); blood pressure (BP); Body-mass-index (BMI); Males (M), Females (F); coronary artery disease (CAD), coronary disease (CD), acute myocardial infarction (AMI), stable angina (SA), unstable angina (UA), Non-obstructive coronary atherosclerosis (NOCA), normal coronary artery (NCA).

**S2 Table. Expression matrix of the most reported molecular entities in CAD.** PRO: protein, MIR: miRNA, MET: metabolite. CluSO identifiers from PADB. Log2(fold-change) and p-value <0.05.

